# Polaramycin B, and not physical interaction, is the signal that rewires fungal metabolism in the Streptomyces – Aspergillus interaction

**DOI:** 10.1101/2022.05.04.490618

**Authors:** Harald Berger, Markus Bacher, Roman Labuda, Isabel Maria Eppel, Florentina Bayer, Michael Sulyok, Erika Gasparotto, Franz Zehetbauer, Maria Doppler, Hannes Gratzl, Joseph Strauss

## Abstract

Co-culturing the bacterium *Streptomyces rapamycinicus* and the ascomycete *Aspergillus nidulans* has previously been shown to trigger the production of orsellinic acid (ORS) and its derivates in the fungal cells. Based on these studies it was assumed that direct physical contact is a prerequisite for the metabolic reaction that involves a fungal amino acid starvation response and activating chromatin modifications at the biosynthetic gene cluster (BGC). Here we show that not physical contact, but a guanidine containing macrolide, named polaramycin B, triggers the response. The substance is produced constitutively by the bacterium and provokes the production of ORS above a certain concentration. In addition, several other secondary metabolites were induced by polaramycin B. Our genome-wide transcriptome analysis showed that polaramycin B treatment causes down-regulation of fungal genes necessary for membrane stability, general metabolism and growth. A compensatory genetic response can be observed in the fungus that included up-regulation of BGCs and genes necessary for ribosome biogenesis, translation and membrane stability. Our work discovered a novel chemical communication, in which the antifungal bacterial metabolite polaramycin B leads to the production of antibacterial defence chemicals and to the up-regulation of genes necessary to compensate for the cellular damage caused by polaramycin B.

## Introduction

Saprophytic fungi are dominant recyclers of natural organic substances specialized in breaking down recalcitrant plant material by their highly effective extracellular enzyme system. However, competition for nutrients, water and space by other organisms such as bacteria may urge fungi to produce defence or signalling compounds, for example antibiotics or other toxic compounds, collectively known as secondary metabolites (SMs), that supress competitors (De Boer et al., 2005; Rousk and Bååth, 2007; de Menezes et al., 2017). The same principle is used by pathogenic fungi which use both extracellular proteins and an arsenal of toxins to penetrate host cells and supress defence mechanisms, respectively (Doehlemann et al., 2017). Apart from their toxic properties, some of these substances are also known to serve as signalling molecules, e.g. for fungal reproduction (Rodriguez-Urra et al., 2012), establishment and maintenance of endophytic or symbiotic associations (Chujo et al., 2019) or pathogenesis-related effectors (Boenisch and Schäfer, 2011; Pusztahelyi et al., 2015). Each fungal species contains genes coding for the biosynthesis of many different SMs but at a given environmental or developmental condition only a specialized set of these secreted molecules is produced. This condition-specific metabolic signature is regulated by genetic signalling networks that have evolved as energy-saving modules for these demanding biosynthetic processes (Yu and Keller, 2005; Gacek and Strauss, 2012; Macheleidt et al., 2016). In many cases the signal triggering transcriptional activation of the biosynthetic genes is connected to nutrient starvation or the onset of reproductive development (Tag et al., 2000; Nemeth et al., 2016). But there are other examples in which rich nutrient sources and active growth is correlated with the production of SMs, such as for the production of siderophores that are required for iron acquisition and metabolism (Oberegger et al., 2001).

The biosynthetic pathways for these metabolites usually encompass many different enzymatic steps and it is a striking feature that biosynthetic and regulatory genes as well as genes encoding transport proteins of a given SM are usually clustered in the genome (termed “biosynthetic gene clusters”, BGCs) and transcriptionally co-regulated. Co-regulation is achieved by cis-acting elements targeted by pathway-specific and broad-domain transcription factors, as well as by chromatin level regulation which facilitates or inhibits access of these regulatory complexes to the target region (Keller et al., 2005; Strauss and Reyes-Dominguez, 2011; Brakhage, 2013). One of the main conclusions drawn from the analyses of the ever-increasing fungal genome databases is the discrepancy between known metabolites and the number of predicted BGCs. For each of the sequenced species, usually only a few SMs are known but many more BGCs are predicted in their genomes (Li et al., 2016). This is not only true for less-studied species but even in aspergilli or penicillia only 20%-30% of the theoretically existing and predicted chemical diversity is known. Taken together this indicates that the majority of BGCs are not expressed under the usually employed standard laboratory growth conditions and we still are not able to mimic the full diversity of growth and developmental conditions that would trigger the production of the full array of metabolites in any of the studied organisms.

One of the triggers that received high attention recently is biotic interactions. It is known since a long time e.g. for pathogenic fungi like *Fusarium graminearum*, that a high proportion of predicted BGCs is only expressed during pathogenic growth (Desjardins et al., 1993; Boenisch and Schäfer, 2011). The corresponding transcriptional program responds to the encountered plant material (substrate response) and to the plant defence system (active plant response). Each of these conditions activates a different subset of SMs in *F. graminearum* (Boedi et al., 2016). There are numerous examples in which biotic interactions of a pathogenic, symbiotic or endophytic life style alters the secondary metabolome of the fungal interaction partner. Also, free-living interactions have been found to alter the SM profiles, e.g. in the mutual interaction between the soil inhabitants *Aspergillus nidulans* and *Streptomyces rapamycinicus* (Schroeckh et al., 2009). In this case, *A. nidulans* produces orsellinic acid (ORS) and yellow polyketides (YPKs), SMs that are barely found in axenic cultures under standard laboratory growth condition (Sanchez et al., 2010). The bacterial trigger ultimately leads to an activating chromatin landscape at the corresponding ORS/YPK BGC (Nützmann et al., 2011) and the formation of this open chromatin structure depends on the function of BasR, a fungal transcription factor induced by amino acid starvation (Fischer et al., 2018b). Consistent with a chromatin-level regulation of the ORS/YPK cluster the same activation can be observed in mutants lacking CclA or KdmB, central components of complexes which regulate the methylation status of histone H3 lysine 4 (Bok et al., 2009; Gacek-Matthews et al., 2016). Also, the addition of the non-metabolizable histidine analogue 3-aminotriazole (3-AT) leads to ORS/YPK induction indicating that the intimate fungal-bacterial contact is not mandatory but bacterial cells or their metabolites may function via an amino acid starvation response and so-far uncharacterized downstream signalling cascades that eventually lead to BasR-mediated chromatin activation (Fischer et al., 2018). How BasR function is connected to the histone H3K4 methylation/demethylation machinery remains to be uncovered.

Using a particular experimental set-up, we identified a specific bacterial macrolide, known as polaramycin B, that is able to trigger ORS/YPK production in *A. nidulans* also in the absence of bacterial cells. We performed RNA-sequencing to test whether polaramycin B application is equivalent to the direct physical contact between the fungal and bacterial cells and found that it fully mimics the transcriptional and metabolic response of the organismic interaction. Strikingly, only Streptomyces species carrying the predicted BGC for guanidine-containing macrolides are able to induce the fungal response.

## Results

### The supernatant of *S. rapamycinicus* is sufficient to induce yellow pigment and orsellinic acid production

To better understand the interaction between *A. nidulans* and *S. rapamycinicus*, we set up an experimental system that allowed us to differentiate fungal hyphae directly interacting with *S. rapamycinicus* from non-directly interacting fungal cells. This setup used fungal and bacterial cultures grown on solid surface in contrast to the previously applied conditions of co-culturing them in liquid shake flasks. In the actual approach the fungal and the bacterial colonies were inoculated at opposing sides of a Petri-dish that contained only agar without nutrients. To allow for some limited growth of the inoculated fungal and bacterial cells, the points of inoculation were supplied with nutrients. This set-up produced a thin, low-density fungal colony spreading out over the agar surface and this allowed us to easily observe colour or morphological changes of the colonies. Figure 1A top shows an example of the experiment in which fungal inoculation points were fed with AMM and bacterial inoculation points with M79 media. Starting from these nutrient reservoirs at the opposite sides of the Petri dish the organisms were able to grow into the nutrient-free agar. Using this setup, we observed that fungal mycelium spreading from at the opposite side towards the bacterial colony was producing some yellow pigments (YPs) although there was no direct contact yet with the *S. rapamycinicus* colony (Figure 1A). No colour changes at the *A. nidulans* mycelia were observed in control experiments growing the fungal cells alone. These observations indicated that yellow polyketides and orsellinic acid may be produced by *A. nidulans* cells in response to the presence of *S. rapamycinicus* already before their physical contact takes place. To make sure there is no direct interaction via buried bacterial or fungal cell interactions we used a similar setup but containing *S. rapamycinicus* cells within a dialysis tube. The molecular weight cut off 8.000-10.000 kDa allowed nutrients passing through the membrane at the point of inoculation but prevented bacterial cells to spread on or through the agar. We found also in this setup an induction of YPs (Figure 1B). To test if the signal for fungal pigment production requires actively growing bacterial cells or if a bacterial metabolite may be sufficient to induce YPs we subsequently applied sterile-filtered culture supernatant from a *S. rapamycinicus* liquid culture grown in M79 media without *A. nidulans*. When adding these bacterial culture supernatants to an *A. nidulans* colony growing alone on the nutrient-free agar we also observed YP production, although colouration was less intense (Figure 1C). Since in this latter approach *S. rapamycinicus* was grown “solo” without *A. nidulans*, we could conclude that production of the YPs in *A. nidulans* is triggered by one or more compounds secreted by *S. rapamycinicus* cells growing in M79 medium and that the production of this inducer(s) does not depend on the presence of a competing or interacting organism. However, the presence of a competing organism like *A. nidulans* may cause elevated production of this unknown molecule(s).

**Figure 1:**
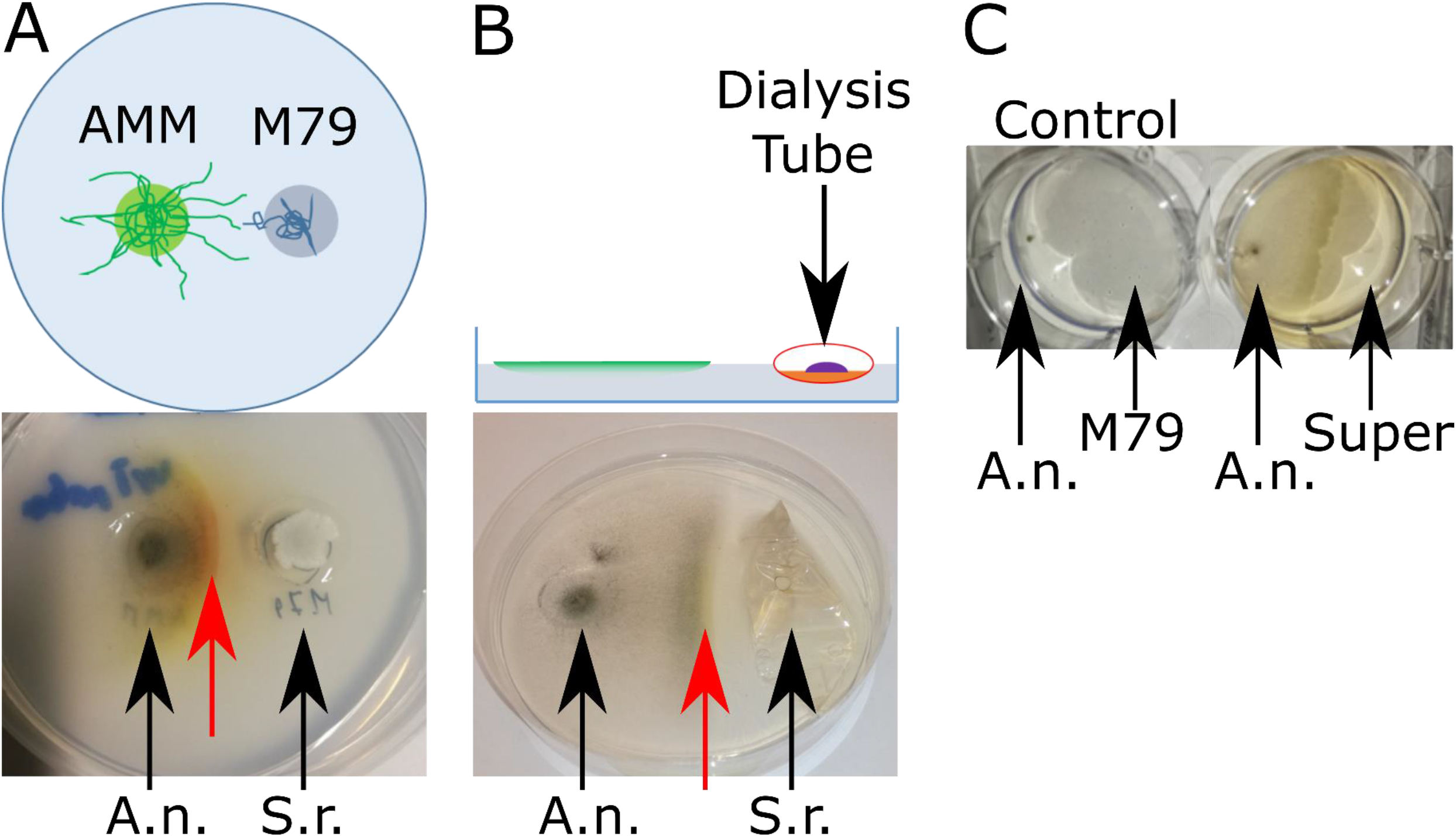
**A**: Top: Schematic setup; Aspergillus minimal medium (AMM) and M79 medium was placed into phosphate buffered water agar (green and blue circles, respectively). AMM and M79 media were inoculated with *A. nidulans* (A.n.) and *S. rapamycinicus* (S.r.), respectively, and incubated for 7 days at 30°C. Bottom: Yellow pigments (YPs) production, (red arrow) of *A. nidulans* growing on phosphate buffered water agar towards *S. rapamycinicus* colonies. **B**: Top: Schematic setup; similar as Figure 1A, except that *S. rapamycinicus* was inoculated into a dialysis tube partially filled with M79 medium thus preventing physical contact with *A. nidulans* cells from the confronting fungal colony. Bottom: red arrow points to the YPs that can also be observed in this setup after 7 days at 30°C, although to a lesser extent. **C**: 10μl of media supernatant (Super) from *S. rapamycinicus* grown “solo” in liquid M79 for 7 days at 30°C, was applied opposite to a growing *A. nidulans* colony (6 well plate) and a clear YPs production was observed.

### Yellow pigment (YP) production correlates with metabolites from the orsellinic acid pathway

As the direct physical *A. nidulans* - *S. rapamycinicus* interaction was shown to trigger the production of ORS and YPs, we then tested if production of the pigments in our test system indeed correlate with these metabolites. For chemical analysis a similar, but larger size set-up of our nutrient-free agar cultivation assay was used (petri dishes with 15 cm diameter) where we grew the organisms for 7 days. Subsequently, in a 9 × 4 grid, agar blocks of 1 cm x 1 cm size were cut out, extracted and metabolites within these agar-blocks were subsequently analysed via photometric absorption measurements at 400nm and by HPLC-MS (Figure 2A and supplementary Figure S1).

**Figure 2.**
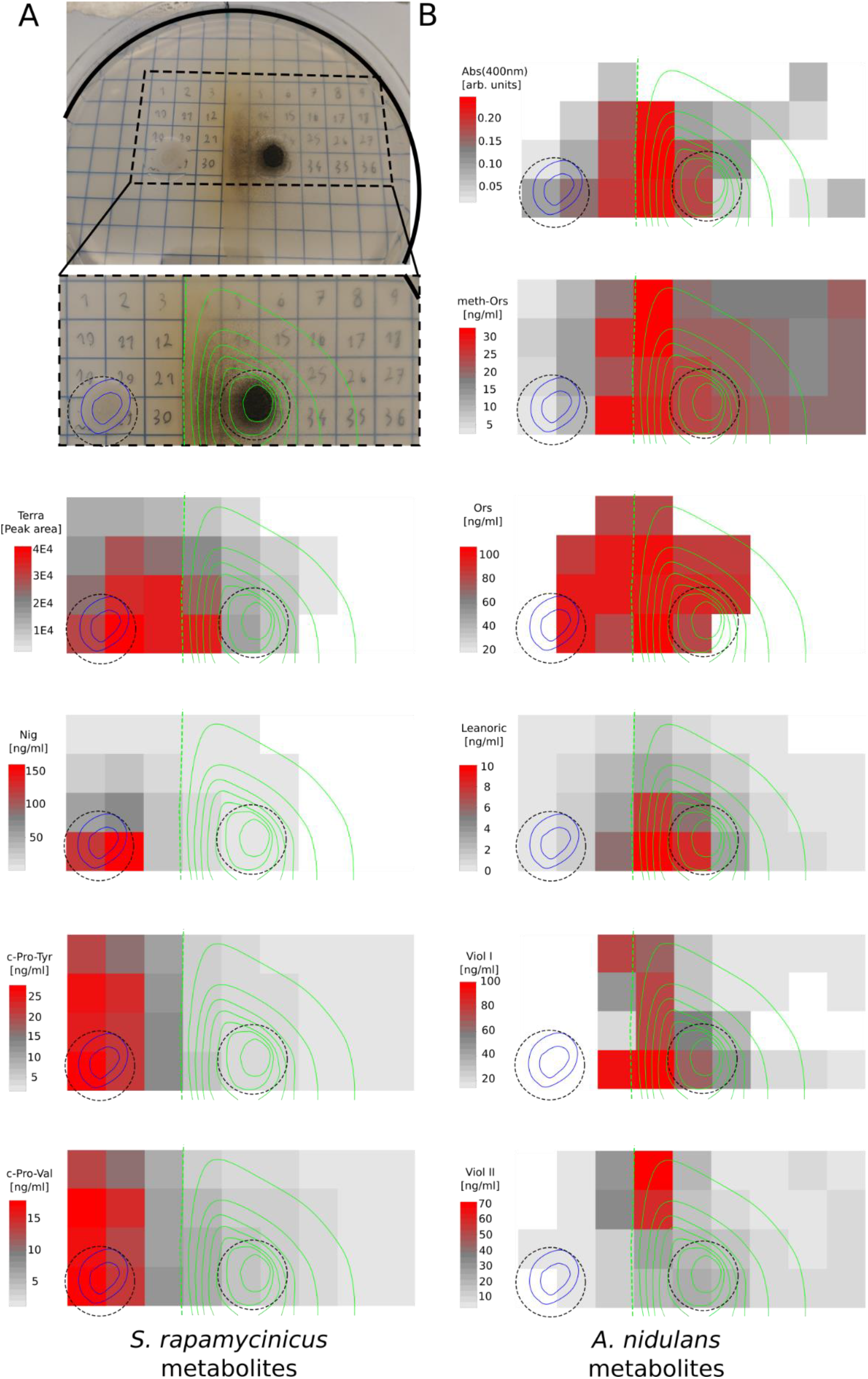
**A:** Top/Left: Experimental setup with an example of the confrontation assay on agar plates and a magnified section of it below the plate photograph. The raster with numbers represent areas that were cut out from the agar and analyzed by mass spectrometry (see materials and methods). Growing *S. rapamycinicus* cells are indicated by blue line circled areas, the *A. nidulans* colony is indicated by green line circled areas; dotted green line shows the extent of fungal growth (complete right hand side of plate is covered by fine mycelia. Dotted black circles indicate the areas of AMM or M79 media in the plate. Conidiation is strongest in the region facing the *S. rapamycinicus* colony. Heatmaps show metabolite concentrations found in the respective regions, positions of colonies are indicated by blue and green lines. Below the photograph in panel A, heatmaps are shown that represent metabolite concentrations found in the respective region originating from *S. rapamycinicus*. **B:** Heatmaps representing metabolite concentrations of the *A. nidulans* colony. Absorption (400nm) representing YPs. Levels of other *A.nidulans* metabolites are shown as separate heatmeaps. Abbreviations: Terra: terragine: Ors: orsellinic acid; Lecanoric: lecanoric acid; meth-Ors: methylorsellinic acid; Viol I: violaceol I; Viol II: violaceol II; Nig: nigericin.

Results of this chemical analyses showed that several metabolites of the ORS biosynthetic pathway were detectable in the extracts obtained from agar samples derived directly from, or from the vicinity of the *A. nidulans* cells growing towards the *S. rapamycinicus* colony. The concentrations of ORS as well as lecanoric acid (LEC) showed strong correlation with the absorption at 400nm that visually appears as yellow colour (Spearmen rho: 0.72 and 0.70 for ORS or LEC, respectively, with p-values of 6.36E-7 or 1.52E-6, see also correlation plot in supplementary Figure S2). These data indicated that our macroscopically observed YPs actually contain ORS and LEC. Additional fungal metabolites have been found in the analyzed agar blocks (Figure 1S). Some of them are also found in higher quantities in the competition zone between fungal and bacterial colonies, but others are more abundant in blocks excised from regions outside of the competition zone.

Interestingly, also *S. rapamycinicus* metabolites seem to be induced by the presence of the fungal mycelium, e.g. terragine is clearly more abundant in the competition zone between the two organisms than anywhere else in the agar of the petri dish. On the other hand, production of the fungal diketopiperazines cyclo(L-Pro-L-Tyr) and cyclo(L-Pro-L-Val) are apparently repressed by the interaction and they show a comparatively lower abundance in this zone (see Figure 2 and supplementary Figure S1).

Taken together, our experiments showed that an active communication based on secreted metabolites takes place between these two organisms and that the cellular responses do not dependent on physical interaction among them. Different concentrations of specific secondary metabolites at different locations within the whole growth area moreover indicate that fungal and bacterial metabolites can be induced or repressed by the chemical signals generated by the cultures.

### The inducer of YPs comprising orsellinic and lecanoric acid is a guanidine-containing polyhydroxyl macrolide

To identify the metabolite(s) from *S. rapamycinicus* inducing YPs in *A. nidulans* we grew *S. rapamycinicus* in the absence of fungal cells “solo” in in M79 medium for 7 days and used a sterile-filtered culture supernatant for the development of an *A. nidulans* bioassay that would monitor YP induction. The fungal bioassay was based on miniature cultures in a 24 well microtiter plate (MTP) format where we grew *A. nidulans* colonies in submerged cultures and were able to use simple colour change of the medium as an indicator of YP production (see details in methods and materials). Using this assay we could confirm that the supernatant of *S. rapamycinicus* cultures were indeed inducing YP in *A. nidulans* and thus can be used for bioassay-guided fractionation tests. Subsequently, several liquid-liquid extraction methods (n-Hexan, Dichlormethan, Acetonitril) were tested and we found that the YP inducer always remained in the aqueous phase indicating that it is a polar substance that may be extractable with methanol. Finally, *S. rapamycinicus* was grown massively on M79 agar plates and the methanol extracts were then further fractionated successively using flash column chromatography and preparative high performance liquid chromatography (HPLC). All fractions were then tested in the bioassay for YP-inducing activity. A highly pure fraction was obtained and purified by preparative HPLC. Subsequent structural elucidation of the active fraction by HPLC-MS/MS and NMR identified the guanidine-containing polyhydroxyl macrolide polaramycin B as the YP, i.e. ORS and LEC inducing compound.

### Polaramycin B biosynthetic gene cluster prediction

To bioinformatically identify the so far uncharacterized polaramycin biosynthetic genes, we performed antiSMASH (Medema et al., 2011) searches of 6 different Streptomyces strains including *S. rapamycinicus, S. lateritius, S. zaomyceticus, S. ederensis, S. europaeiscabiei* and *S. iraniensis* based on published genome sequences and annotations. As polaramycin is similar to azalomycin, a well-characterized guanidine-containing macrolide, we assumed that similarity to the known azalomycin biosynthetic gene cluster (BGC) would be indicative for a BGC potentially responsible for polaramycin production. Only in *S. rapamycinicus* and *S. iraniensis* a homology with the azalomycin F3a biosynthetic gene cluster from Streptomyces sp. 211726 (MIBiG accession: BGC0001523) could be detected (Figure 4). To substantiate the hypothesis, we also cultivated isolates of four additional Streptomyces species under identical conditions as done before for *S. rapamycinicus*. Strikingly, only the two species *S. rapamycinicus* and *S. iranensis*, which contained the predicted gene cluster similar to the azalomycin cluster in their genomes, induced YP production. In contrast, the other four species not harbouring these BGCs showed no YPs (see supplementary Figure S3). These findings suggest that the region showing homology to the azalomycin biosynthesis cluster is responsible for the production of polaramycin in *S. rapamycinicus* and *S. iraniensis* and thus for YP induction.

**Figure 3.**
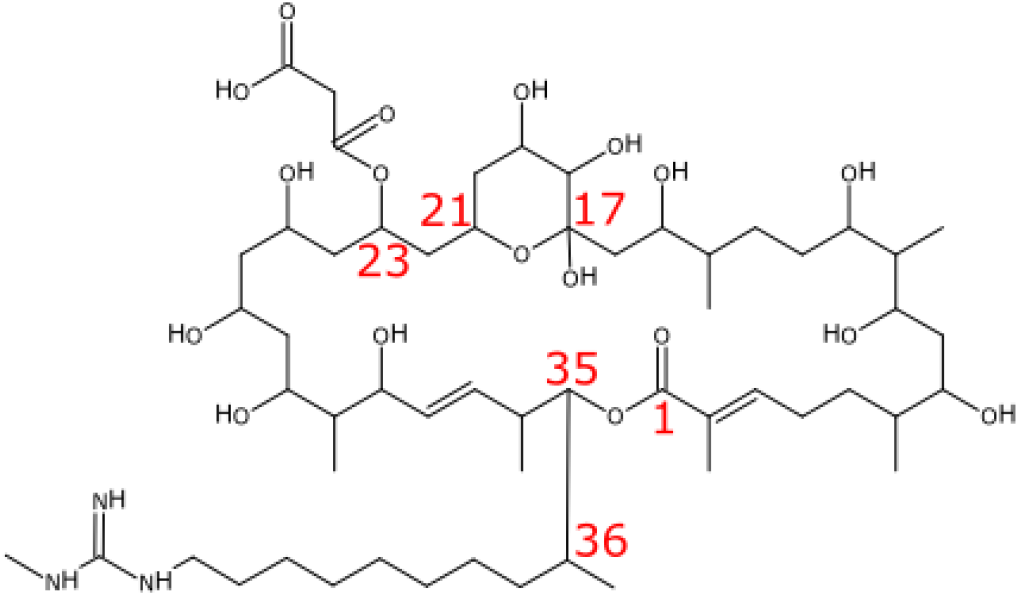
Structure of PolaramycinB. The C17-C21 hemiketal ring has been shown to be important for antimicrobial activity, whereas C23 malonyl reduced antimicrobial activity.

**Figure 4.**
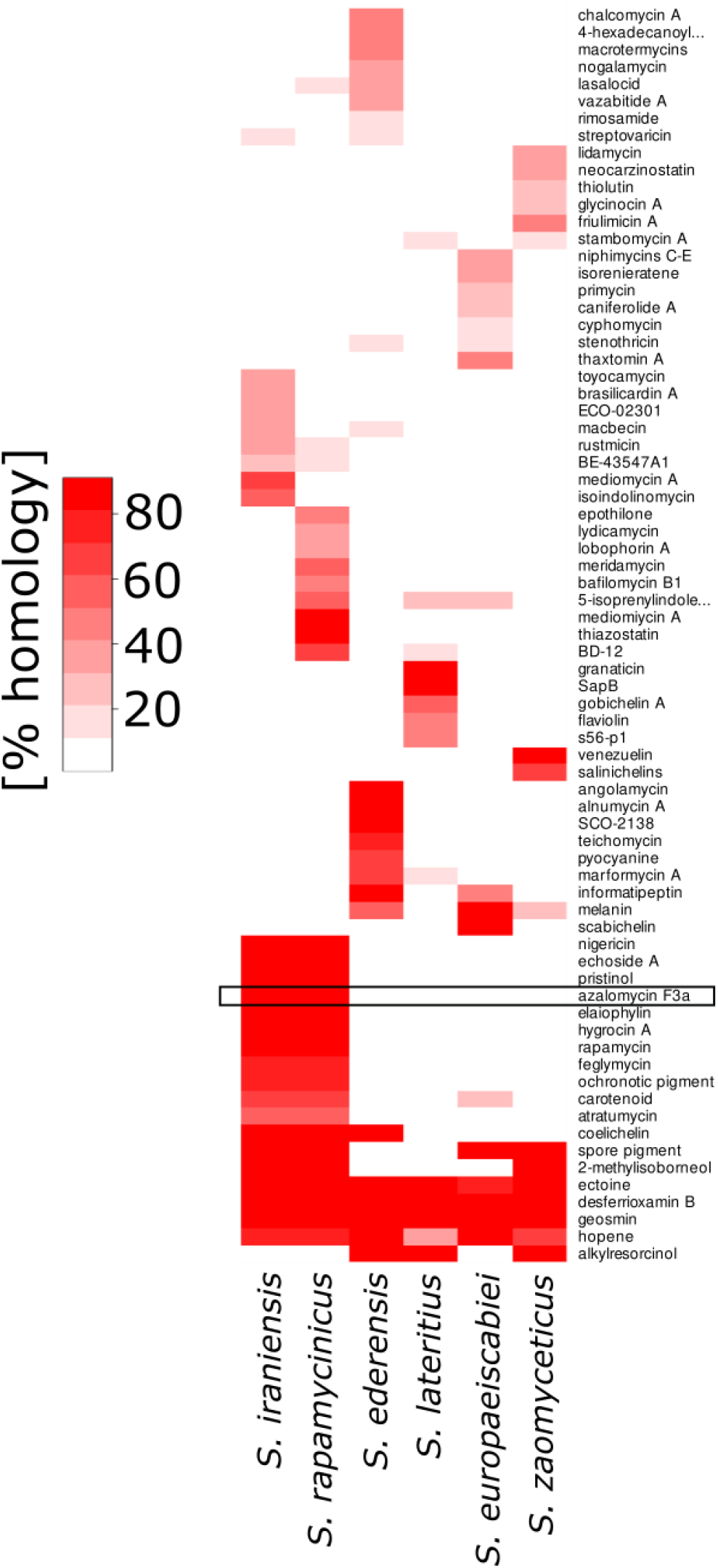
Homology search (anti-SMASH) in *S. rapamycinicus, S. iraniensis, S. zaomyceticus, S. ederensis, S. lateritius* and *S. europaeiscabiei* for putative secondary metabolite clusters. Only *S. rapamycinicus* and *S. iraniensis* induce YPs, both containing a homologous azalomycin F3a gene cluster (black box). Red color indicates percent homology.

**Figure 5:**
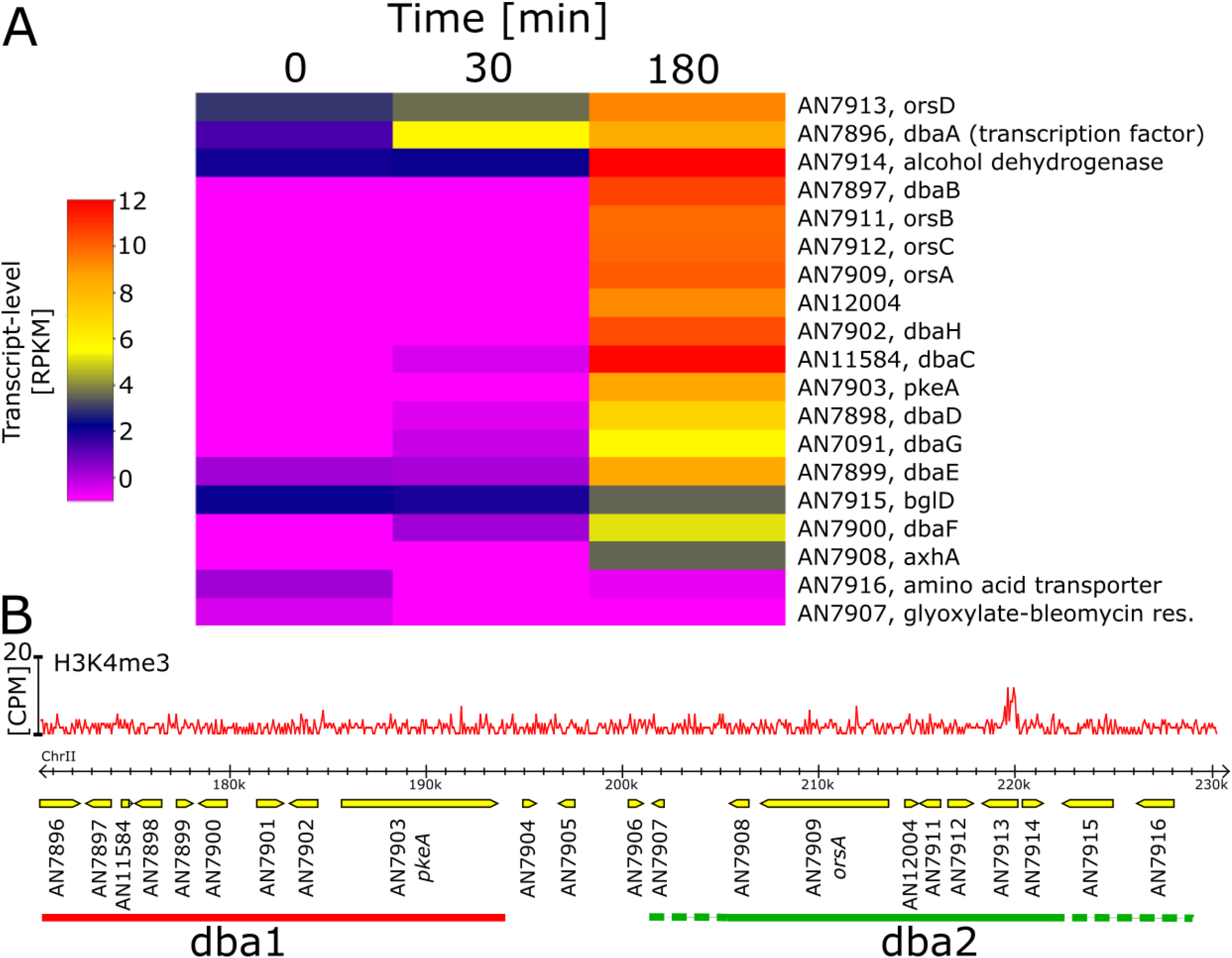
**A**. Color coded transcript levels (RPKM) of dba cluster genes, columns represent time after polaramycin B treatment. Gene names or descriptions are shown if available. The transcription factor *dbaA* is already induced after 30 minutes following treatment, what seems not to be the case for the remaining cluster genes, that are induced at the later time point, presumably only after the transcription factor protein DbaA was translated and activated. AN7916 and AN7907 are not induced at all and may not be part of the dba-cluster. **B:** Genome and chromatin organization of both dba1 and dba2 derivative of Benzaldehyde1 gene clusters. Red graph shows histone H3K4 methylation after 18 hours, taken from (Gacek-Matthews *et al.*, 2016), a single significant peak found in AN7913.

### Dose Response of polaramycin B

Since induction of orsellinic acid production was previously only observed when *S. rapamycinicus* and *A. nidulans* interacted physically, we reasoned that in these former liquid shake co-cultures the local concentration of polaramycin B at the fungal cell surface may not have been sufficiently high to induce the metabolic changes. In this former set-up, the concentration necessary to trigger ORS induction may only have been reached when cells directly interacted with each other because this physical interaction prevented immediate diffusion of the inducer away from the fungal cell surface. We therefore tested YPs production and *orsA* gene expression in *A. nidulans* cultures treated with different concentrations of polaramycin B in our 24 well plate bioassay. We found that after 48 hours of incubation a concentration of 0.5 μg/mL polaramycin B in the *A. nidulans* growth medium is necessary to induce YPs as well as *orsA* transcription (Figure_S4). This concentration was then used in our genome-wide transcriptional analysis aimed at determining whether polaramycin B treatment induced a similar genetic response as previously observed in the physical interaction and if other fungal secondary metabolites are also affected by the addition of polaramycin B treatment. Moreover, the identification of transcriptional network changes by the treatment is of interest to understand which fungal targets are affected by this compound.

### Polaramycin B induces the dba1-dba2 gene cluster for ORS biosynthesis

For RNA-seq analysis, we used *A. nidulans* cultures pre-grown for 18 hours in AMM shake cultures and subsequently added polaramycin B at a concentration of 0.5 μg/ml. As the transcriptional network changes triggered by the substance may differ depending on the duration of treatment, we chose two different harvesting times. To test for the short-term response, we harvested the mycelia after 30 minutes of incubation and for the response to an extended treatment time the cultures were harvested after 3 hours of incubation with polaramycin B. It needs to be mentioned that at this time point there is still 0.45− 0.5% glucose remaining in the AMM medium, a concentration that is known to repress starvation-induced secondary metabolites, such as the production of sterigmatocystin. To control if under these conditions polaramycin B also induces YPs and ORS, we first tested the expression levels of *orsA* (AN7909), the polyketide synthase-encoding gene. Using RT-qPCR we could confirm the induction event (data not shown) and we thus used these samples for RNA sequencing via Illumina Hi-Seq.

In respect to the ORS gene cluster our data analysis showed that the genes of the split dba1-dba2 cluster are induced by polaramycin B treatment. Moreover, the regulatory gene *dbaA*, which encodes the specific transcription factor of this BGC, and which resides within the dba1 cluster, is induced already after 30 minutes. Consistent with DbaA being the cluster-specific transcriptional activator protein, other genes within the dba1-dba2 BGC, were found to be induced after three hours post-treatment. Interestingly, one gene of unknown function (AN7913, designated *orsD)* and located within the dba2 region also shows the same induction kinetics as the transcription factor (induction after 30 minutes). Other dba1 cluster genes residing in the neighbourhood of the *dbaA* transcription factor gene also seemed to be induced after 30 minutes. This result that may hint towards a local chromatin effect that renders the *dbaA*-containing region transcriptionally active.

### Polaramycin B also induces other BGCs

Our transcriptome data allowed to analyse whether other BGCs were also affected by the polaramycin B treatment. Indeed, we found that several other BGCs are also changed in their transcriptional activity. For example, a majority of genes in the terrequinone (Bouhired et al., 2007) or aspercryptin (Henke et al., 2016) BGCs are significantly upregulated. Some BGCs showed only partial activity, such as the asperfuranone or the AN10289 cluster (Klejnstrup et al., 2012), in which only roughly half of the predicted cluster genes are induced. Interestingly the penicillin cluster, whose 3 genes are already transcribed at intermediate levels in our overnight cultures, are significantly downregulated by polaramycin B treatment (see Table 1). Thus, apart from the YPs and few other substances, the majority of BGCs do not show significant alterations in their expression profiles. This transcriptional response is similar to what has been also observed in the previous experiments in which *A. nidulans* cells experienced direct physical contact with *S. rapamycinicus* (Schroeckh et al., 2009). These findings indicate that polaramycin B is the responsible substance that mediates the transcriptional response of A.nidulans in the *S. rapamycinicus* interaction.

**Table 1:** Selection of BGCs with differentially regulated genes (between 0 hour and 0.5 hour or between 0.5 hour and 3 hour samples). Up-regulated, Down-regulated and total refers to number of genes at least 2-fold up- or downregulated and total number of genes predicted or shown to belong the given cluster, respectively.

| BGC | Up-regulated | Down-regulated | total |
| --- | --- | --- | --- |
| Derivative of Benzaldehyde1 (dba) and F9775 hybrid cluster 1 & 2 | 17 | 0 | 19 |
| AN7884 (Aspercryptin)cluster | 13 | 0 | 15 |
| Terrequinone (tdi) cluster | 4 | 0 | 5 |
| Penicillin cluster | 0 | 2 | 3 |
| AN5318 cluster | 2 | 1 | 5 |
| Asperfuranone (afo) cluster | 4 | 1 | 9 |
| AN9129 cluster | 1 | 0 | 2 |
| AN9314 cluster | 1 | 0 | 2 |
| pkdA cluster | 5 | 0 | 11 |
| AN10289 cluster | 2 | 0 | 5 |

### The induction kinetic suggests possible modes of action of polaramycin B

We have been involved in the elucidation of the BasR and chromatin-based mechanisms by which *S. rapamycinicus* changes the *A. nidulans* metabolism during their physical interaction (Fischer et al., 2018a). To identify putative targets of polaramycin and analyse the possible overlap between physical interaction and chemically triggered responses, the transcriptional profiles were grouped into 8 clusters (see Figure 6): Immediately upregulated to steady state (A1); Immediately downregulated to steady state (A2); Immediately upregulated and continuously further upregulated (B1); Immediately downregulated and continuously further downregulated (B2); Delayed upregulated (C1); Delayed downregulated (C2); transiently upregulated (E1); transiently downregulated (E2).

**Figure 6.**
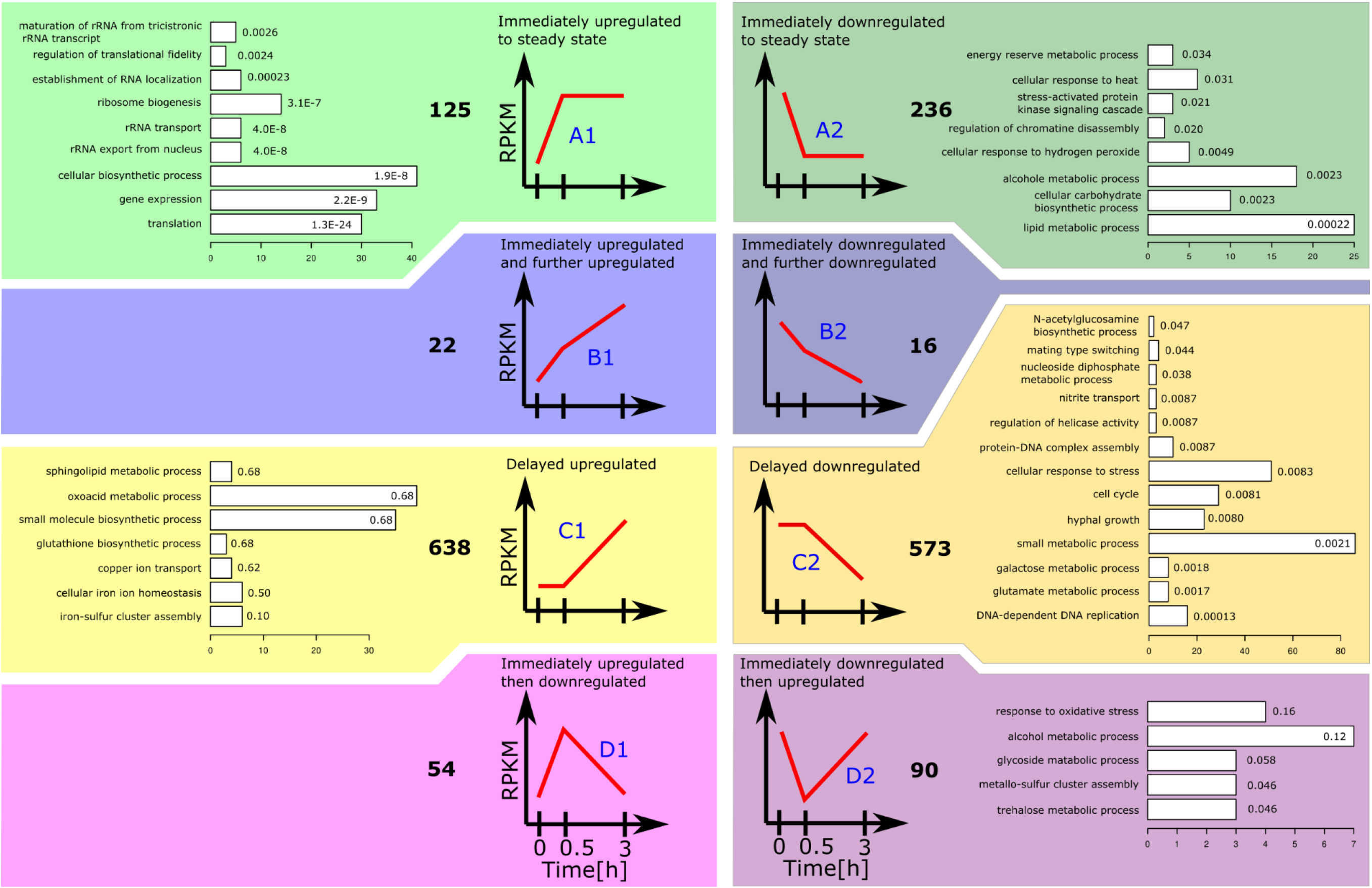
Gene-set clustering of induction kinetiks in response to Polaramycin B at 3 time points. The combination of a p-value < 0.01 and a differential expression of ±1 was condidered a significant differential expression. y-axses of diagramms show transcript levels, x-axes represent time. Total number of genes in each clustered gene-set are shown in bold numbers, barplots show GO-term enrichments, bar-length represent number of genes, number within barplots show corrected p-values.

A significant regulation was assumed if absolute log2 difference was greater than 1 and p-value was less than 0.01. The contrasts considered were between time point zero and 30 minutes as well as between 30 minutes and 3 hours. In case of disagreement between log2 difference and p-value no differential regulation was assumed. To identify processes possibly involved in the polaramycin response we searched for overrepresented gene ontology (GO) annotations in each gene-set (Ashburner et al., 2000).

### Genes immediately responding to polaramycin B treatment

Interestingly, in the gene-set of genes “immediately upregulated to steady state “ (A1, 151 genes) GO annotations associated with translation or ribosome biosynthesis were significantly overrepresented. The genes with these annotations in cluster A1 were already transcribed at intermediate levels but polaramycin B caused a significant increase at least 2 fold.

In addition to the translation process, also few genes putatively associated with membrane stability and function are found in this immediately induced gene set like AN3125 (predicted role in response to stress and integral to membrane localization), AN7327 (predicted glycosylphosphatidylinositol GPI-anchored protein), AN9342 (predicted role in transmembrane transport and integral to membrane localization) among the highest differentially expressed genes.

In the gene-set of genes “immediately downregulated to steady state” (A2, 236 genes) the GO annotation “Lipid metabolic process” was significantly enriched which supports the findings that the antifungal activity of macrolides (like Azalomycin F) is based on disturbance and leakage of the phospholipid cell membrane (Sugawara, 1967). But in our experiments we found only few genes differentially regulated with the GO annotation “Response to osmotic stress” (e.g. AN065) which could be a hint that the cell membrane disturbance caused by polaramycin B has no large impact or can be well counteracted by the fungal metabolism, such as by those induced genes mentioned above coding for membrane stability functions.

The rather small gene-set B1 of genes “immediately and continuously upregulated” (22 genes) mainly contains the upregulated BGCs, such as 2 genes from the dba-cluster and 3 genes from the AN7884-cluster (aspercryptin) as well as one from the AN0016-cluster (*pes1*). Statistically, no significant GO-term enrichment could be detected what may be due to the small size of this gene-set.

Also the gene-set B2 of genes “immediately and continuously downregulated” is rather small (16 genes) and contains mainly transport-related genes. For example, AN9168 (putative solute-hydrogen symporter activity with role in glycerol transmembrane transport), AN6277 (predicted role in transmembrane transport), AN4277 (putative glucose transmembrane transporter activity), AN3489 (putative endoplasmic reticulum localization) and AN6071 (domains with predicted membrane localization) are found in this gene set.

### Genes only responding to extended polaramycin B incubation

We could observe that a larger number of genes is only differentially expressed after prolonged treatment with polaramycin B. These genes were grouped into sets C1 and C2 comprising of 638 upregulated and 573 downregulated genes. The upregulated gene-set C1 showed no concise picture in respect to enriched GO-terms but next to a diverse array of predicted gene functions, it contained basically all upregulated BGCs discussed above. Most of the dba-cluster genes fall into this C1 gene-set as well as several other secondary metabolite cluster genes(see Table 1) In addition to BGCs, some of the strongest induced genes have a linkage to oxidative stress responses, like AN7893, which has predicted oxidoreductase activity, AN9315 has a role in NADH oxidation and regulation of reactive oxygen species, AN5397 is a copper-containing laccase that oxidizes phenolic substrates or AN2559 with predicted nucleotide binding and oxidoreductase activity. The fact that these genes are upregulated upon prolonged polaramycin B treatment may be a result of a disturbed membrane potential triggered by the substance eventually causing oxidative stress. If this stress is the “real” inducer of YPs remains to be determined.

Predicted functions of genes downregulated upon prolonged polaramycin treatment (gene set C2) are associated with GO terms like hyphal growth, DNA replication or cell cycle regulation. This indicates that polaramycin B treatment impairs growth processes, a result that we have already observed in our phenotyping plate assays (Figure 1 and 2).

As can be seen in Figure 6 for the gene-set C2 general catabolism related GO terms fall into this category. These include AN7180 and AN7541, which are two cutinases involved in carbohydrate catabolism, AN11188 with a putative role in endocrocin biosynthetic process, AN2719 with cell wall peptidoglycan catabolic process, AN11159 with phospholipid biosynthetic process. Also, many proteolytic or hydrolytic enzymes are found in this gene-set: AN7121 (predicted metallocarboxypeptidase), AN5558 (alkaline protease), AN111981 (predicted UDP-N-acetylmuramate dehydrogenase), AN1583 and AN0009 (predicted hydrolase), AN3393 (similarity to neutral metalloprotease II), AN7962 (deuterolysin-type metallo-proteinase), AN8445 (Putative aminopeptidase). The complete list of genes is shown in supplementary Table ST2. The downregulation of these hydrolytic enzymes may be a result of the reduced growth probably to avoid self-digestion or a reduced turnover of metabolites to not allow potential competitors to feed easily on such generated metabolites.

### Genes with a transient response to polaramycin treatment

There are genes that have a strong short-term response to the treatment with polaramycin B (measured after 30 minutes exposure) but this response is lost upon prolonged treatment with the drug (measured after 3 hours exposure) Gene-set D1 holds 54 genes which are immediately upregulated by the presence of polaramycin B but no response is observed any more in the 3 hours treatment cultures. No clear picture comes out from this gene set but 2 GPI- anchored proteins (AN7792 a Putative lysophosphoplipase A and AN8609), AN2798 (predicted role in response to stress and integral to membrane localization), AN0209 (mepB, High-affinity ammonium transporter), AN4372 (*pgaB*, a protein with polygalacturonase activity) have been found in this gene-set. This may hint towards a reaction on the membrane due to polaramycin B that is transient as the cell may be able to balance the membrane defects after the respective genes have been induced and new membrane components have been produced.

Also, quite a significant number of genes were found to be transiently repressed by polaramycin B treatment. 90 genes were found in this set of genes designated D2 (Figure 6). Interestingly 3 genes associated with trehalose, an important storage carbohydrate in fungi are in this gene-set, AN9340 (*treA*), AN10533 and AN5523 (*tpsA*). Since trehalose plays also an important role in conidial stability, this may be a reflection of the dynamics following the immediate downregulation of carbohydrate metabolic processes (see gene-set A2) post polaramycin B treatment

## Discussion

### Polaramycin B structure, function and concentration thresholds for changing fungal metabolism

The molecular details of the interaction between microorganisms that compete for physical space and/or metabolic resources is still a relative sparsely investigated field of molecular microbiology despite its omnipresence in our biosphere. In previous work it was suggested that a direct, physical contact between *S. rapamycinicus* and *A. nidulans* is necessary to trigger the genetic response of the fungus leading to the production of orsellinic acid (ORS) and its derivates (Schroeckh et al., 2009b). Using a novel experimental system, we were now able to identify the chemical molecule that triggers the genetic response underlying the ORS production in these co-cultures. We identified a 36 carbon-membered polyol macrolide containing a hemiacetal monoester of malonic acid and a guanidyl group, collectively known as polaramycin B. This substance has previously been extracted from *Streptomyces hygroscopicus* LP-93 but to the best of our knowledge not much research has been done on this particular variant of guanidine-containing polyhydroxyl macrolide since its discovery in 1997 (Meng and Jin, 1997).

Generally these macrolides have been shown to possess broad-spectrum antibacterial and antifungal activities (reviewed in (Song et al., 2019)). By comparison of different variants of these macrolides, the following assumptions about structure-function relationships could be made: The size of the lactone ring has minor influence on the antimicrobial activity (32 and 36 sized structures were compared), but the terminal guanidine group is crucial for their antibacterial and antifungal activities. The six-membered hemiketal ring (C17-C21) plays an essential role in the antimicrobial activity, the malonyl group (C23) reduces the antimicrobial activity and reduction of double bond or methylation of the lactone ring does not have a strong influence on the antimicrobial activity (see (Song et al., 2019) and references therein).

Most likely, in many natural environments the concentration of polaramycin B will not reach or exceed the threshold concentration of 0.5μg polaramycin B per ml of medium, as determined in the artificial laboratory growth condition. Therefore, in nature, direct interaction between bacterial and fungal cells will most likely still be required for effective chemical communication, simply because the triggering molecules freely diffuse. And this may also have been the reason why in high-volume liquid shake co-cultures the required concentration of polaramycin B was only reached when cells directly interacted. Also, the fungal cell density plays a role as higher cell densities dilute the number of polaramycin B molecules per fungal cell. We conclude this because in densely growing *A. nidulans* cultures incubated on high-nutrient AMM culture plates (55 mM glucose, 10mM nitrate) the YPs-inducing effect by opposing *S. rapamycinicus* colonies was very hard to observe compared to the water-agar medium lacking these nutrients. In the latter, YPs were easily detectable in the very fine and thin mycelial network that *A. nidulans* cells develop under these conditions.

### The metabolic response of fungal colonies to bacterial confrontation is unidirectional

The water-agar plate assays allowed us to observe the metabolic “warfare” between the two microorganisms and we could observe that the fungal colony produces YPs and ORS only in the region that faces the bacterial colony, but not in the far side and in distant areas of this competition region. That indicates that the fungal cells respond individually to the presence of the bacterium and that this signal is not transferred very far through the mycelial network. So, a fungal colony only seems to represent a limited “information network” that does not lead to a genetic alert of cells not directly connected to each other. Macroscopically, a fungal colony appears as an interwoven network of cells, however, most likely at the cellular level, they may not be truly connected as one would expect distant cells of a colony also responding to the presence of polaramycin B in a similar way the cells facing the confrontation zone, do. Apart from the observed production of YPs and other secondary metabolites, this spatially limited response was also observed for other phenotypic changes caused by polaramycin B, such as a lack of growth inhibition and onset of sporulation at the distant, far side of the competition zone. The conidiation phenotype was corroborated with data from our transcriptome analyses in which we observed highly transcribed sporulation related genes (e.g. AN3624 (*ygA*), AN4998(*gapA*), AN2513 (*pipA*), AN0928, AN7735 (*eglD*) and AN0863 after polaramycin B treatment. This observation indicates that a fungal colony, although representing a network of connected cells, does not represent an “information network” in the sense that a signal arriving at one end of the colony is passed on to the other side of the same colony.

### Possible mode of action of polaramycin B

Beside the dba1-dba2 cluster for ORS and YPK production, also other BGCs showed increased transcription levels in response to polaramycin B. These included aspercryptin and the terrequinone clusters, but due to lacking metabolite standards, missing mass-spectrometry data or low metabolite concentrations the clear-cut correlation between transcriptional levels and metabolite concentrations cannot be made at the moment. On the other hand, not all *A. nidulans* BGCs seem to respond equally to polaramycin B treatment. For example, the penicillin cluster in is repressed by this substance. This is interesting in light of the fact that Streptomycetes strains are known producers of penicillin and similar antibiotics. Therefore, these bacteria require a strong self-resistance mechanism against these types of antibiotics and apparently, *A. nidulans* actively ceases production of an antibiotic that is obviously useless against this specific competitor (reviewed in (Ogawara, 2016)).

Previously, it has been shown that a closely related molecule to polaramycin B, azalomycin F, acts on fungi via disturbance of the cell membrane, but not much work was committed into deciphering its mode of action. We applied transcriptomics to gain more insight and could identify genes related to lipid metabolism, membrane function and growth regulation as being down-regulated during the treatment. Apparently, affected cells responded to this challenge by up-regulation of possible compensatory genetic networks, such as genes necessary for membrane stability, for translation and ribosomal functions as well as enzymes that counteract oxidative stress. This response overlaps with the 3-Amino-triazol (3-AT) treatment and is also similar to the response of the direct physical interaction between *A. nidulans* and *S. rapamycinicus*, (Nützmann et al., 2011; Fischer et al., 2018a). 3-AT exposure is known to block histidine biosynthesis eventually leading to an amino acid starvation and induction of the cross-pathway control genes like *cpcA* and *basR* (Braus et al., 2006) Our groups also found chromatin mutants with changes in the ORS and YPs production.

### A chromatin-related function of polaramycin B in activating BGCs?

Chromatin structure is influenced by histone modifications and in fungi, transcriptional co-regulation of the genes residing inside a certain BGC, is facilitated by chromatin transitions (Strauss and Reyes-Dominguez, 2011; Pfannenstiel and Keller, 2019). Interestingly, a mutant lacking a component of the COMPASS complex that is responsible for activating histone H3K4 methylation (H3K4me3) was found to strongly overproduce ORS and YPKs and also featured strongly induced expression of the dba1-dba2 cluster (Bok et al., 2009). Consistently, the opposite type of mutant lacking the H3K4me3 de-methylase gene KdmB, showed lower expression of these BGCs (Gacek-Matthews et al., 2016; Bachleitner et al., 2019). As H3K4me3 is a histone modification generally associated with activating functions, it seems counter-intuitive that a mutant lacking this function would be more active in transcription of the ORS/dba cluster. However, it is known that cells lacking the H3K4 methyltransferase system have a defect in silencing loci positioned at or near chromosome (Nislow et al., 1997; Margaritis et al., 2012). Additionally, or alternatively, the lack of H3K4 methylation may downregulate a basic metabolic gene involved in amino acid biosynthesis thus leading to amino acid starvation and subsequent induction of ORS/YPs production. The experimental proof of this hypothesis awaits further research.

## Materials and Methods

### Strains and cultivation conditions for transcriptomics

Aspergillus nidulans WT (pabaA1 veA1) was the fungal strain used throughout the study (Pontecorvo et al., 1953). *Streptomyces rapamycinicus* strain was the one used by Fischer *et* al. in our laboratory for the transcriptome and chromatin analysis (Fischer et al., 2018). Conidiospores from cryostocks were plated on Aspergillus minimal medium (AMM) plates (10mM nitrate, 1% Glucose, 15 pM para-amino benzoeic acid), incubated for three days at 37 °C and harvested in autoclaved deionized water (diH2O) containing 0.1 % Tween-20. A spore density of 1E6 spores/mL was inoculated in 50 mL of liquid AMM in 100 mL Erlenmeyer flasks, respectively, and the culture grown at 37 °C and 180 rpm. The time of cultivation was 18 hours. Treatment with polaramycin B (0.5μg/ml) for 30 minutes and 3 hours or no treatment was applied.

### Starvation plate assay

Starvation plates were agar plates containing only AMM salts without nutrients (0.52g/l KCl, 0.815g/l KH2PO4, 1.045g/l K2HPO4), 2 circular regions (diameter: 15 mm) were cut out and filled with AMM- or M79-agar and inoculated with 1 μl of Aspergillus or Streptomyces spore suspensions (5E6 and 5E8 spores /ml), respectively. For metabolite analysis 1×1×0.5 cm agar blocks were cut out and metabolites were extracted in 3 ml Acetonitril-Acetic acid-Water (79:1:20) solvent.

### Bioassay

The bioassay to detect yellow pigment production in *A. nidulans* was designed for 24 well plates. 1E+6 *A. nidulans* spores were incubated per ml AMM and grown in submerse culture in 50 ml AMM at 30°C shaking (180rpm) flask overnight. Single fungal pellets were placed in wells of a 24 well plate and 2ml of AMM media not containing trace elements was added. The plate was then incubated for 2-3 days at 37°C. No sporulation could be observed till this time points. These plates were then used for testing extracts.

### Streptomyces fermentation

Liquid fermentation of *Streptomyces rapamycinicus* was performed in M79 medium using 1E+7 spores/ml in 50 ml at 30°C 160rpm. Solid state fermentation was performed on M79 agar plates (10g/l Glucose, 10g/l bacto-peptone, 1g/l casamino acids, 2 g/l yeast extract, 6 g/l NaCl, pH 7.2). For the final polaramycin B preparation 2000ml M79-agar was distributed into 40 petri-dishes (large) and each one inoculated with 1E+7 spores/plate. After 7 days of growth at 30°C agar was cut into approximately 1cm x 1cm pieces and extracted with 1.5 L methanol over night at room temperature. The filtered methanol fraction was then evaporated to a volume of 5ml.

### Isolation of the macrolide

The crude extract was purified by reversed-phase silica gel vacuum flash chromatography (Interchim, puriFlash^®^450), using three consecutive Interchim puriFlash^®^ 32 g silica IR-50C18-F0025 flash columns (particle size: 50μm). The columns were eluted with a binary solvent gradient (solvent A: H2O, solvent B: ACN). The starting linear gradient from 10% B to 27% B during 25 min at a flow rate of 15 mL/min was followed by an isocratic elution at 52% B for 10 min. Then a linear gradient from 52% to 66% B over 7min was applied at the same flow rate and finally the column was washed starting with 100% B for 10 min followed by 100% A for 10 min at a flow rate 15-30 mL/min. UV 254 nm and UV scan 200-400 nm modes were used for detection and final separation of 8 main fractions (F1-F8), which were consequently concentrated under reduced pressure at 45°C. The target compound (activity) was found in fraction F5 (18-20 Rt, yield: 25 mg).

This fraction was further dissolved in 1:1:1; ACN/CH3OH/H2O and purified by an Agilent 1260 Infinity preparative HPLC (USA) on a reversed phase column Gemini NX C-18 (21.20 x 150 mm, 5 μm, 110 Å). Gradient starting with 30 % ACN and 70 % H2O up to 95 % ACN in 32min (total time 45 min) and a flow rate of 25 ml/min. Four preparative fractions (pF1 -pF4, bioactivity guided fractionation, time lapse fractionation) were collected, of which pF3 was found as active. The active fraction was further proceeded with 2nd stage prep HPLC (signal lapse fractionation). The active signal (target) was found at tR 10.5 min, yielding 0.91mg. For purity check an Agilent 1200 system was used with the same stationary phase and gradient program.

### Secondary metabolite and Orsellinic acid quantification

Metabolite analysis was carried out using a 1290 Series HPLC System (Agilent, Waldbronn, Germany) coupled to a QTrap 5500 LC-MS/MS System (Applied Biosystems SCIEX, Foster City, CA) equipped with Turbo Ion Spray electrospray ionization source as described earlier (Sulyok et al., 2020). Chromatographic separation was performed at 25 oC on a Gemini^®^ C18-column, 150 × 4.6 mm i.d., 5 μm particle size, equipped with a C18 4 x 3mm i.d. security guard cartridge (Phenomenex, Torrance, CA, US). 5μl of sterile filtered sample were directly injected without any further manipulation.

Confirmation of positive metabolite identification was carried out by the acquisition of two time scheduled multiple reaction monitoring (MRMs) which yielded 4.0 identification points according to the European Commission decision 2002/657. In addition, retention time and ion ratio had to agree to the related values of authentic standards within 0.03 min and 30% rel., respectively. Quantitation was based on external calibration using serial dilutions of a multi-analyte stock solution. The accuracy of the method is verified on a continuous basis by participation in a proficiency testing scheme organized by BIPEA (Gennevilliers, France) with a current rate of z-scores between −2 and 2 of > 94 % (> 1300 results submitted).

### RNA extraction and expression analysis

RNA was extracted from liquid nitrogen frozen mycelium using TRIzol Reagent (Thermo Fisher Scientific). 2 μg total RNA were then subjected to DNase I digestion (Thermo Fisher Scientific). cDNA was amplified using iScript™ cDNA Synthesis Kit (Bio-Rad) and LunaScript^®^ RT SuperMix Kit (NEB). cDNA was diluted 1:10 and used for qRT-PCR (Bio-Rad CFX384).

### RNA sequencing and analysis

Total RNA extraction samples were transferred to Vienna Biocenter Core Facilities (https://www.viennabiocenter.org/vbcf/) for library preparation and Illumina high throughput sequencing using poly-A enrichment kit (NEB) and Nextera Library prep kit. 50 bp single end sequencing was performed using a HiSeq v4 Illumina sequencer. Obtained sequences were de-multiplexed, quality controlled, filtered using trimmomatic 0.36 (Bolger et al., 2014) and mapped on the *Aspergillus nidulans* genome assembly (A_nidulans_FGSC_A4_version_s10-m03-r07). Mapping was performed using BWA (Li and Durbin, 2009) and reverse transcripts were counted using python script HTSeq (Anders et al., 2015). Normalization and statistics were done using R/Bioconductor and the limma and edgeR packages, using mean-variance weighting (voom) and TMM normalisation (Law et al., 2014). A significance cut-off of p < 0.01 was applied for analysis. Transcription levels are log2 read counts per kilobase of exon per million library reads (RPKM). For trace graphs transcript coverage was calculated as explained for the ChIP-seq experiments to obtain counts per million reads (CPM). All data are available at NCBI GEO under the accession number GSE185285.

### GO-enrichment analysis

To simplify GO interpretation we used semantic similarity measures to identify major ontologies enriched in specific experiments. This means we clustered the observed GO terms based on their distance in the acyclic GO-graph (Sayols et al., 2016) and extracted a common parent term.

### NMR- measurement

All NMR spectra were recorded on a Bruker Avance II 400 (resonance frequencies 400.13 MHz for 1H and 100.63 MHz for 13C) equipped with a N2-cooled 5 mm broadband cryoprobe head (Prodigy) with z–gradients at room temperature with standard Bruker pulse programmes. The sample was dissolved in 0.6 ml of MeOD (99.8 % D, Eurisotop, Saint-Aubin, France). Chemical shifts are given in ppm, referenced to residual solvent signals (3.31 ppm for 1H, 49.0 ppm for 13C). 1H NMR spectrum was collected with 32k complex data points whereas the 13C-jmod spectrum with WALTZ16 1H decoupling was acquired using 64k data points. All two-dimensional experiments were performed with 1k × 256 data points, while the number of transients and the sweep widths were optimized individually. For the TOCSY experiment the spin-lock time was set to 100 ms and the spinlock field to 8.3 kHz. HSQC experiment was acquired using adiabatic pulse for inversion of 13C and GARP-sequence for broadband 13C-decoupling, optimized for 1J(CH) = 145 Hz.

## Acknowledgements

We are grateful to Axel Brakhage and his lab members for provision of bacterial strains and critical discussion of the manuscript. Work was supported by grants from the OEAD “Sparkling Science” program (grant N° SPA 06-258) and from the Austrian Science Fund FWF, grant N° P32790-B.

