## Supplementary Figures for "Polaramycin B, and not physical interaction, is the signal that rewires fungal metabolism in the Streptomyces – Aspergillus interaction"

Supplementary Figure 1

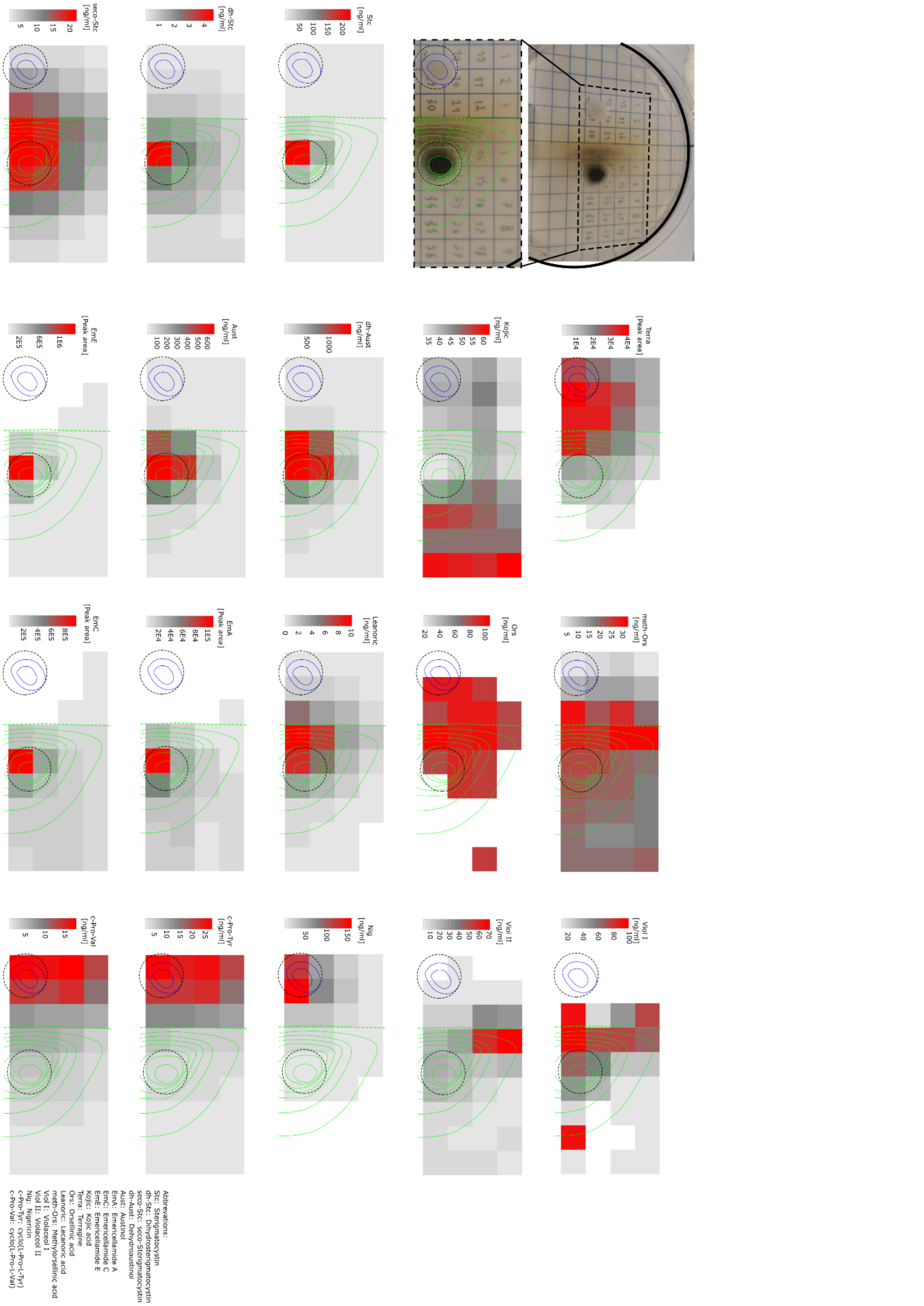


Top/Left: Experimental setup, raster with numbers were cut out and analyzed. Growing *S. rapamycinicus* is indicated by blue edging, *A. nidulans* colony is indicated by green edging; dotted green line shows extent of fungal growth (complete right hand side of plate is covered by fine mycelia. Dotted black circles indicate extent of AMM or M79 media. Conidiation is strongest in region facing the *S. rapamycinicus* colony. Below: heatmaps showing metabolite concentrations found in the respective region originating from *S. rapamycinicus*, positions of colonies are indicated by blue and green lines. Bottom: Heatmaps representing concentrations of Bacterial metabolites found in the respective positions in the agar plate. Respective metabolite concentrations from the *A.nidulans* or *S. rapamycinicus*. Abbr.: Terra: terragine: Ors: orsellinic acid; Lecanoric: lecanoric acid; meth-Ors: methylorsellinic acid; Viol I: violaceol I; Viol II: violaceol II; Nig: nigericin; EmA, EmC, EmE: Emericellamid -A, -C, -E; Stc, dh-Stc, secco-Stc: Sterigmaticystein, dehydro- secco-Sterigmatocystein; Aust, dh-Aust: Austionol, dehydro-Austinol; c-Pro-Tyr: cyclo(L-Pro-L-Tyr); c-Pro-Val: cyclo(L-Pro-L-Val)

Supplementary Figure 2

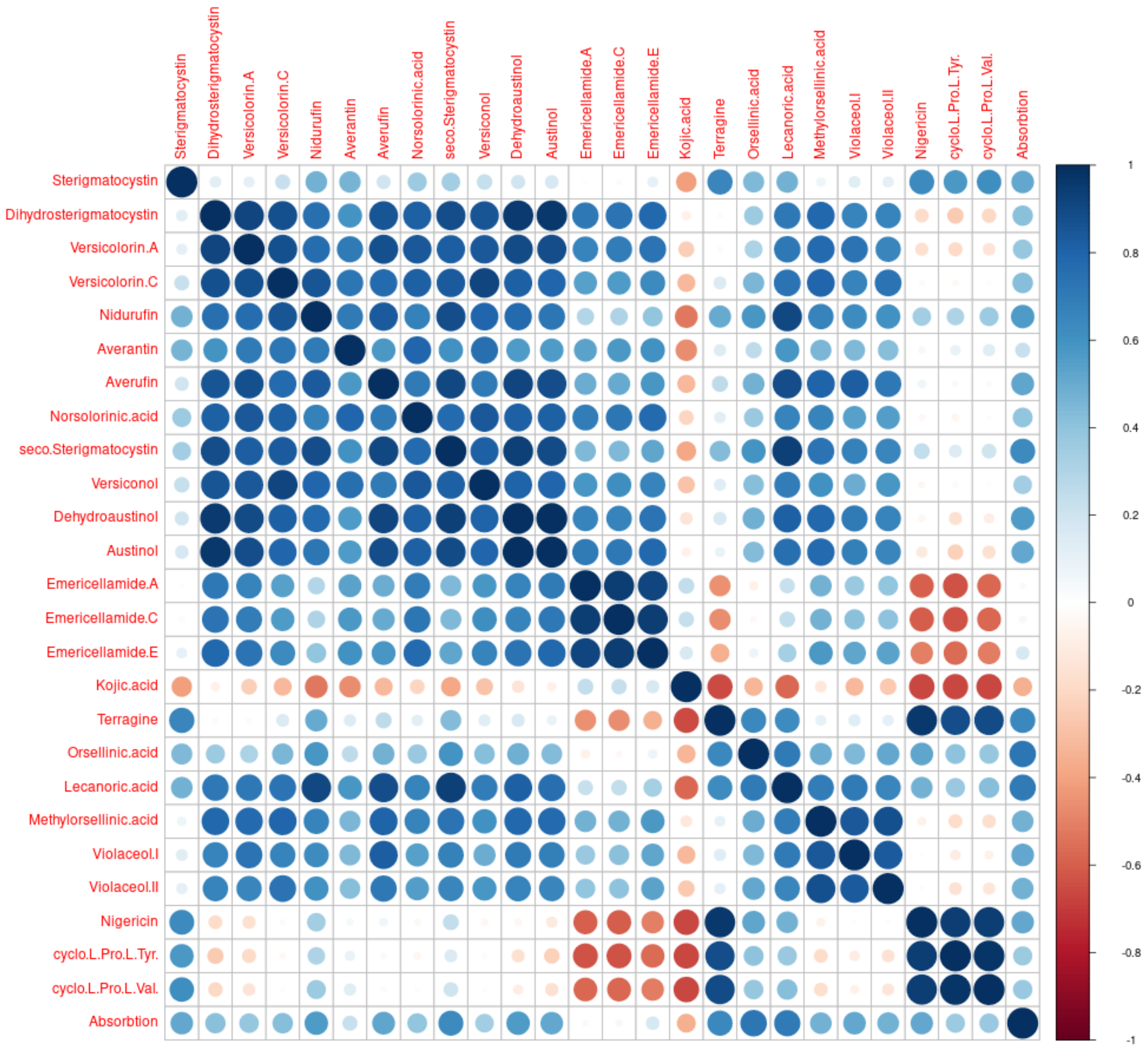


Figure S2: Spearman correlation plot of metabolites in Agar plate. Blue: positive correlation, red: negative correlation.

Supplementary Figure 3

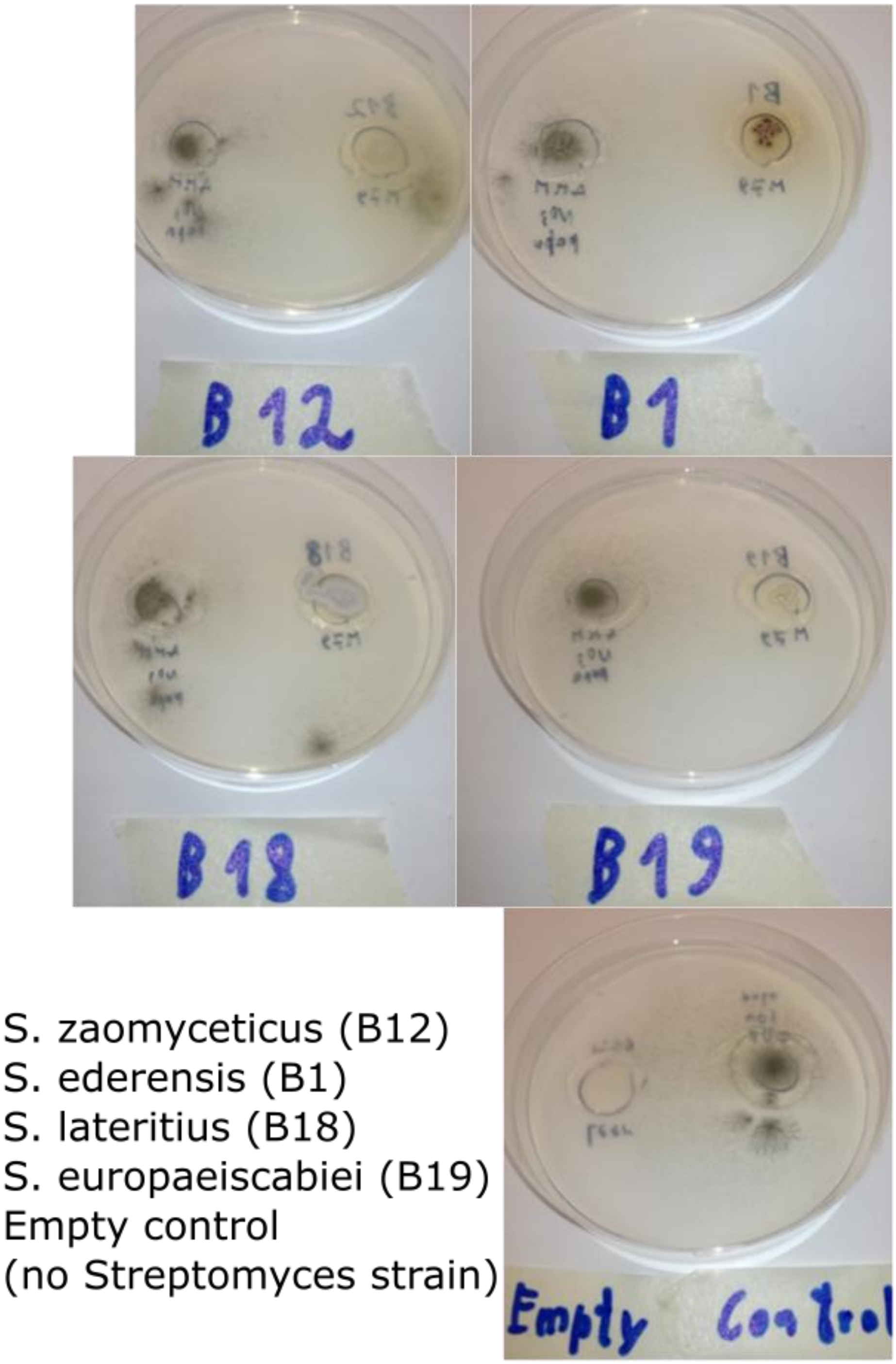


Figure S3: Agar plate test of *S. zaomyceticus*, *S. ederensis*, *S. lateritius* and *S. europaeiscabiei* for inducing yellow pigment production. Neither of these strains contains a homologous azalomycin F3a gene cluster.

Supplementary Figure 4

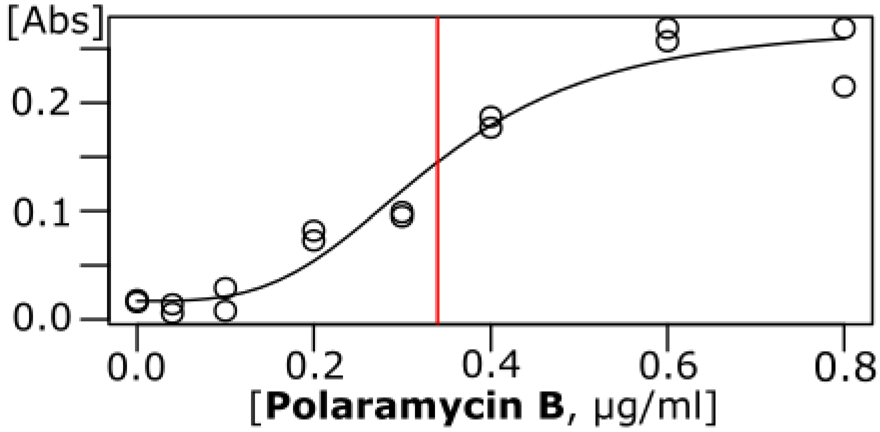


Figure S4: Dose response of Polaramycin B in respect to YP production (absorption at 400 nm) in *A. nidulans*
